# Parental assigned chromosomes for cultivated cacao provides insights into genetic architecture underlying resistance to vascular streak dieback

**DOI:** 10.1101/2024.05.26.595985

**Authors:** Peri A. Tobias, Jacob Downs, Peter Epaina, Gurpreet Singh, Robert F. Park, Richard J. Edwards, Eirene Brugman, Andi Zulkifli, Junaid Muhammad, Agus Purwantara, David I. Guest

## Abstract

Diseases of *Theobroma cacao* disrupt cocoa bean supply and economically impact growers. Vascular streak dieback (VSD), caused by *Ceratobasidium theobromae,* is a new encounter disease of cacao currently contained to southeast Asia and Melanesia. Resistance to VSD has been tested with large progeny trials in Sulawesi, Indonesia, and in Papua New Guinea with the identification of informative quantitative trait loci (QTL). Using a VSD susceptible progeny tree (clone 26), derived from a resistant and susceptible parental cross, we assembled the genome to chromosome-level and discriminated alleles inherited from either resistant or susceptible parents. The parentally phased genomes were annotated for all predicted genes and then specifically for resistance genes of the *nucleotide-binding site leucine-rich repeat* class (NLR). On investigation, we determined the presence of NLR clusters at informative QTLs, as well as other potential disease response gene candidates. Understanding the genetics underlying resistance and susceptibility to VSD will accelerate the breeding cycle by providing clear targets for molecular screening. Additionally, we provide the first diploid, fully scaffolded and parentally phased genome resource for *Theobroma cacao* L.

## Introduction

Cacao, *Theobroma cacao* L. (Malvaceae) is an important perennial tree crop producing beans that are used as raw materials in the chocolate, beverage, and cosmetic industries (Schnell *et al*., 2007). Much of the world’s cocoa bean production comes from small-holder, mixed cropping farms in some fifty countries that depend on trade with developed countries where the raw product is manufactured into chocolate (Daymond *et al*., 2022). Cacao is a difficult crop to grow profitably due to several endemic problems. Alongside other problems, diseases disrupt the supply of cocoa and have serious economic impacts to growers (Marelli *et al*., 2019). Responses to disease can lead to tree removal and replacement with more profitable and reliable crops (Daymond *et al*., 2022). The centre of origin for *T. cacao* is the Upper Amazon, South America, where evolutionary studies indicate divergence from close common ancestors around 9.9 million years ago (Richardson *et al*., 2015). While genetic diversity exists within the geographic centre of origin, cultivation largely relies on a limited genetic pool of *T. cacao* (Motamayor *et al*., 2008). Southeast (SE) Asia is one of the major suppliers of cocoa beans and cultivation has historically been relatively disease-free due to the lack of co-evolved pathogens (Bailey and Meinhardt, 2016). However in the 1960s, vascular streak dieback (VSD) emerged as a new encounter disease of cacao and caused devastating crop losses in Papua New Guinea (PNG) (Guest and Keane, 2007).

The causal agent of VSD is *Ceratobasidium theobromae*, a fastidious biotrophic fungus, first discovered and described by Philip Keane (1972). *Ceratobasidium theobromae* then spread to other SE Asian and Melanesian cacao regions as an emergent disease on cacao (Guest and Keane, 2007). In Indonesia, the pathogen has spread into Sumatera Regions (Trisno *et al*., 2016) and was subsequently confirmed in a new area of Barru District, Sulawesi (Junaid *et al*., 2020). The pathogen is currently contained to the SE Asia region but presents a serious biosecurity threat to cacao in West Africa and Latin America. Symptoms include green spotted chlorosis in older leaves and vascular browning in the petioles and stems that often lead to rapid branch or whole tree death in susceptible genotypes (Guest and Keane, 2007; Samuels *et al*., 2012). Some genotypes of *T. cacao* that were resistant to disease 50 years ago remain resistant, indicating that durable resistance is highly heritable and therefore may be exploited in breeding to introgress resistance into new, improved cacao genotypes (Tan and Tan, 1988). Varieties of cacao that show resilience to infection by *C. theobromae* have been studied in both PNG and Sulawesi, Indonesia. Results of these studies identified informative quantitative trait loci (QTL) that correlate with both qualitative and quantitative resistance in progeny populations (Epaina, 2012; Singh, 2021). The two independent studies used progenies derived from crosses between Trinitario **KA2-101** (resistant to VSD) and **K82** (susceptible to VSD) in PNG (Epaina, 2012), and between Amelonado **S1** (resistant to VSD) with Iquitos, Criollo and Amelonado **RUQ 1347** (susceptible to VSD and formerly known as CCN 51) in Sulawesi, Indonesia (Singh, 2021). The progeny trials used 346 and 130 progeny that segregated in response to VSD in PNG and Sulawesi, respectively. The informative QTLs associated with resistance indicated a likely polygenic response and were mapped onto nine of the ten *T. cacao* chromosomes from the PNG progeny trial (Epaina, 2012) and three in the Sulawesi trial (Singh, 2021). The two studies used either simple sequence repeat (SSR) or single nucleotide polymorphism (SNP) markers and compared linkage maps with established disease resistance mapping developed by Lanaud et al. (2009). The available genomes for Matina 1-6 (Motamayor *et al*., 2013) and Criollo (Argout *et al*., 2011) were used to inform the linkage maps. Interestingly, VSD QTLs co-localized with a region identified for resistance to Phytophthora Pod Rot (PPR) and Frosty Pod Rot (FP) (causal agents *Phytophthora palmivora* and *Moniliophthora roreri*, respectively) on chromosome 8, while others were common for response to both VSD and PPR on chromosomes 3, 4 and 5 in only one study. Both studies identified VSD QTLs within similar regions on chromosomes 3, 8 and 9. These apparent shared QTL regions indicate hotspots for disease resistance.

Classical selection and breeding for resistance, particularly in perennial trees, can be a long and difficult process (Sniezko, 2006). The research is further complicated for resistance to *C. theobromae* as the pathogen cannot be cultured in vitro and is extremely slow growing within the host plant before visible symptoms develop. Traditional Koch’s postulates cannot be satisfied for biotrophic organisms, however molecular approaches are becoming increasingly important in validation studies (Bhunjun *et al*., 2021). Understanding the genetics underlying resistance and susceptibility to VSD is likely to accelerate the breeding cycle by providing clear targets for molecular screening.

Effective plant responses to disease at the molecular level are well established as involving resistance genes of the *<u>n</u>ucleotide-binding site leucine-rich <u>r</u>epeat* family (NLRs) (Fluhr *et al*., 2001). NLR genes are known to be numerous, clustered, and highly polymorphic both between and within species (Barragan and Weigel, 2020; Santos *et al*., 2022; Weyer *et al*., 2019; Van de Weyer *et al*., 2019). We therefore approached the problem of understanding the underlying genetics of resistance to VSD by building on the structural knowledge of NLR genes and from the progeny QTL studies of Epaina (2012) and Singh (2021). We aimed to leverage the latest sequencing and bioinformatic approaches to gain a clearer insight into the genetic regions of interest. Our premise for the work was that (i) QTLs are informative for predicting resistance, (ii) genetic resistance hotspots are present, (iii) QTLs may indicate the presence of NLR gene clusters, (iv) polymorphic NLR genes within clusters, from contrasting genotypes, may explain disease phenotype. The questions we therefore posed were, can we assemble the cacao genome and successfully discriminate alleles inherited from parents of known VSD phenotype within an F1 tree progeny, and if so, can we determine differences in NLR clusters, type, and specific gene sequences at the QTL locations, based on parentage? An additional question that we tested was, using new sequencing technologies, can we assemble and scaffold to chromosome level both pathogen (∼30 Mb) and host (∼380 Mb) genomes from a single sample? This would circumvent the inability to culture the biotrophic *C. theobromae* and provide insight into the genomics of a compatible interaction. This final aim was not successful due to the limited pathogen DNA we were able to extract, but future studies could work to improve on this approach.

Taking these premises and goals, we built a high-quality parental phased genome resource for a susceptible cacao (clone 26) of known parental cross, **S1 maternal** and **RUQ 1347 paternal**. We annotated the genome for all predicted genes, and then specifically for only NLR-type genes, and conducted fine scale investigation of genomic regions associated with resistance to *C. theobromae* to determine the resistance gene complements inherited from both parents. We made comparative studies into the NLR genes, structurally, quantitatively, and qualitatively. Our investigations show structural differences in NLR presence within parental chromosomes and provide insights into resistance to VSD and potentially other serious diseases of cacao. Additionally, we provide the first diploid, fully scaffolded and phased genome resource for *Theobroma cacao* L.

## Results and Discussion

### Near complete parental assigned chromosomes for cultivated *Theobroma cacao*

We assembled and parentally phased the chromosomes of a VSD susceptible *T. cacao* (clone 26), the progeny of a cross between a maternal resistant (S1, haplotype A) with a paternal susceptible (RUQ 1347, haplotype B) tree. All but three inherited chromosomes (two S1, one RUQ 1347) show telomeric caps (black dots in Figure 1). The genomes show few gaps (red plus symbol in Figure 1), high levels of contiguity and completeness, as well as chromosomal synteny (Edwards *et al*., 2022) based on conserved single copy orthologs (Simão *et al*., 2015). Minor inverted translocations on chromosomes four and nine (pink synteny lines) are likely to be biologically real considering the lack of flanking gaps that would indicate mis-scaffolding. The phased genomes are both highly complete according to assessment of conserved single copy orthologs and basic statistics (Table 1).

**Figure 1.**
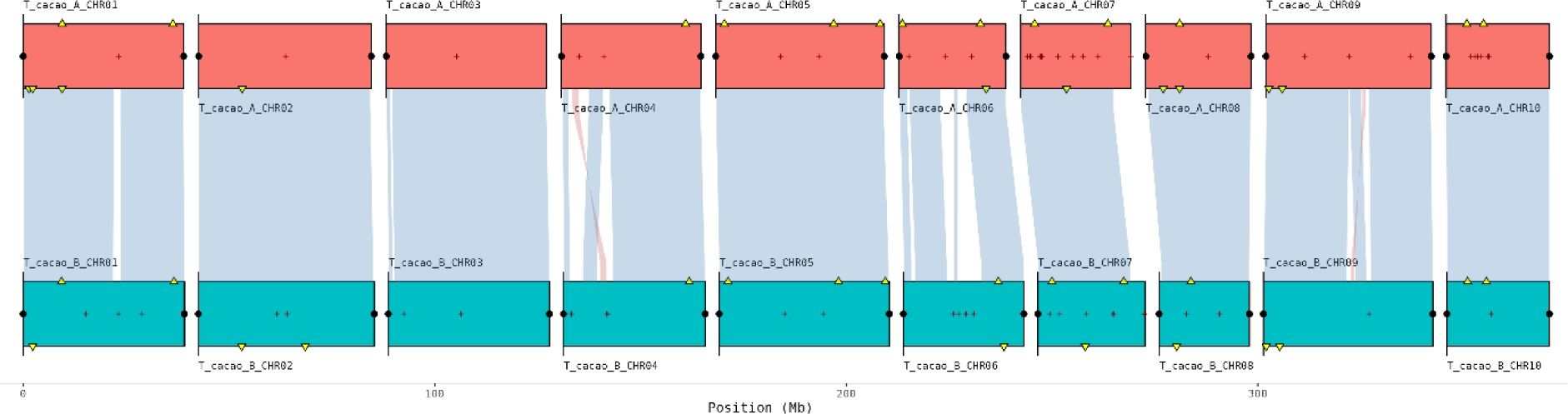
Chromsyn (Edwards *et al*., 2022) synteny plots of Theobroma cacao parentally phased genome, haplotype A (top) and B (below). Synteny blocks of collinear “Complete” BUSCO genes (Simão *et al*., 2015) link scaffolds from adjacent assemblies: blue, same strand; red, inverse strand. Yellow triangles mark “Duplicated” BUSCOs. Filled circles mark telomere predictions from Telociraptor (black) (Edwards, 2023). Assembly gaps are marked as dark red + signs.

**Table 1.** **Raw data statistics from NanoPlot (De Coster and Rademakers, 2023) and genome statistics for both sets of parentally inherited *Theombroma cacao* chromosomes using Quast and BUSCOs** (Gurevich *et al*., 2013; Simão *et al*., 2015). **Haplotype A chromosomes derived from maternal plant S1 and haplotype B from paternal plant RUQ 1347. Annotation statistics based on FindPlantNLRs and Funannotate** (Chen *et al*., 2023; Palmer and Stajich, 2023). **NLR annotated resistance genes were classified and largely conformed to coiled-coil class with only small numbers of Toll/Interleukin-1 receptor-type (TIR) and integrated domain type NLRs.**

| Raw HiFi data |  |  |
| --- | --- | --- |
| Total bases | 30,259,224,851 |  |
| Mean read length | 14,413.4 |  |
| Mean read quality | 35.6 |  |
| Genome statistic | Haplotype A | Haplotype B |
| Total genome length (bp) | 374,523,129 | 381,631,648 |
| chromosome length (bp) | 338,223,858 | 340,140,478 |
| No. of chromosomes | 10 | 10 |
| N50 (bp) | 38,945,508 | 39,156,173 |
| L50 | 5 | 5 |
| No. of contigs | 165 | 315 |
| GC (%) | 33.73 | 33.67 |
| **BUSCO complete (genome; *n* = 1,614)** | **98.2%** | **98.4%** |
| Single-copy | 97.5% | 97.8% |
| Duplicated | 0.7% | 0.6% |
| BUSCO fragmented | 0.6% | 0.9% |
| BUSCO missing | 0.3% | 0.7% |
| **Protein-coding genes (funannotate)** | 26,199 | 26,321 |
| **NBARCs (FindPlantNLRs annotation)** | 342 | 370 |
| NLRs | 293 | 326 |
| TIR-NLR (TNL) | 11 | 15 |
| Integrated domain (ID) NLRs | 5 | 12 |
| **Repeats** | **62.0%** | **62.4%** |

The goal for genome assemblies of gapless and telomere-to-telomere (T2T) chromosomes has recently been achievable with the latest sequencing technologies (Sergey *et al*., 2022). The resulting genome assemblies permit greater depth in understanding structural and evolutionary biology (Mao and Zhang, 2022). Recently, chromosome-level genomes for three wild cacao species from the Upper Amazon were made public (Nousias *et al*., 2024). These provide the first cacao genomes produced using long read sequence technology and indicate an evolutionary divergence, based on conserved orthologs, from cultivated Criollo and Matina reference genomes (Argout *et al*., 2011; Motamayor *et al*., 2013) at ∼ 1.83 and ∼ 1.34 million years ago. The wild cacao assemblies represent the collapsed haploid state for genomes and are larger in both chromosome and genome base pair (bp) lengths compared with earlier reference genomes. Each of our parentally phased genome assemblies are comparable in size to the wild cacao, at around 380 Mb, however we annotated more predicted protein coding genes at around 26,000 (Table 1) compared to around 21,000, and higher percentage of repetitive sequences at 62% compared to 53% (Nousias *et al*., 2024). We annotated all genes for the predicted nucleotide binding site (NBARC) domain type as well as predicted ‘complete’ NLRs that incorporate NBARC plus leucine-rich repeat (LRR) domains (Table 1). Of the complete NLRs we determined only 11 and 15 Toll/Interleukin-1 receptor-type TNL-type, and only 5 and 12 novel integrated domains (ID) within NLRs per haplotype.

### Low structural diversity of NLRs between *Theobroma cacao* alleles

The NLR-type resistance genes were annotated, as described in the experimental procedures, and the parentally inherited chromosomes were compared, noting that haplotype A (red annotations (Figure 2Figure 2) aligned with the resistant maternal and haplotype B (yellow annotations) with susceptible paternal trees. We found that the predicted NLR-type genes varied in quantity and types between the two sets of chromosomes. Of the 342 and 370 predicted genes that contained NBARC domains (Table 1), one of the key identifiers of this class of NLR-type resistance genes, our research determined a discrepancy of 293 and 326 complete gene models within Haplotype A and B respectively. Using the web tool version of OrthoVenn3 (Sun *et al*., 2023) with the OrthoMLC algorithm, using an expect value of 1e-15 and inflation value 1.5, we found that NLR amino acid sequences clustered into 83 orthologous groups (orthogroups). Of these orthogroups, 82 clusters were shared between the two haplotypes with the largest cluster including 55 proteins, 28 and 27 from haplotype A and B respectively. Forty-four orthogroups contained a single NLR protein from each haplotype, meaning that the matching homolog was present in each assembly. Nevertheless, aligned sequences for homologous proteins showed some variations and there were also 2.4% single NLRs, numbering 4 and 11 from haplotype A and B respectively.

**Figure 2.**
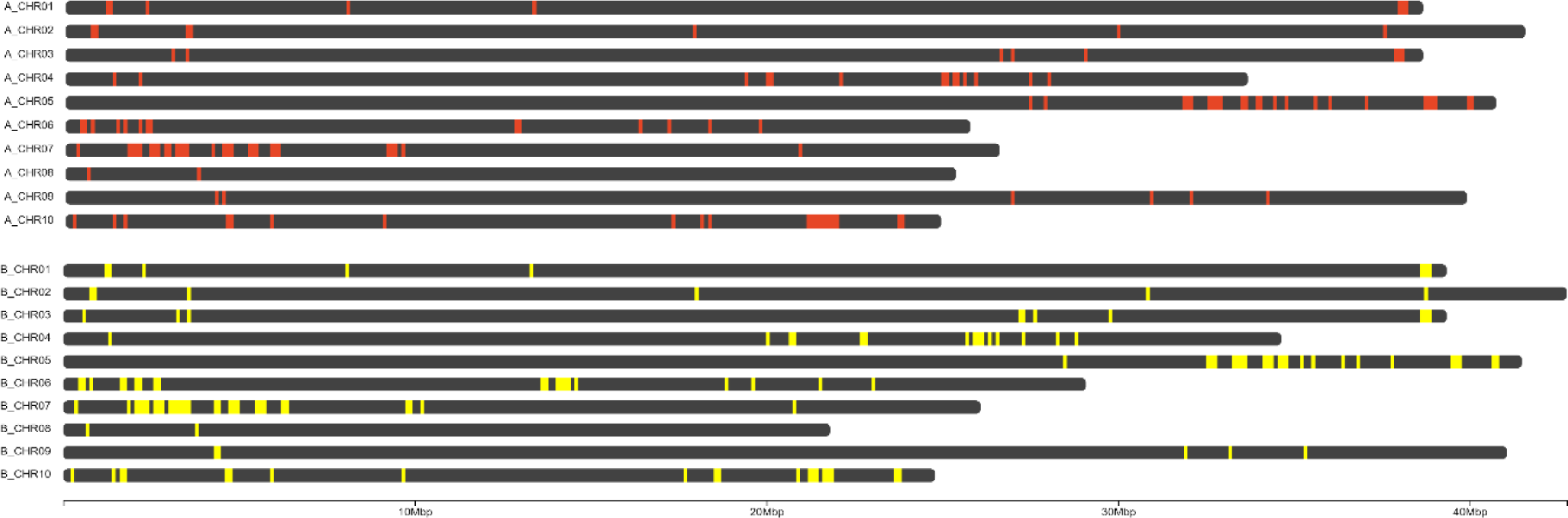
Physical locations of predicted NLR genes on the paired chromosomes of *Theobroma cacao* (A) haplotype A (red) and (B) haplotype B (yellow) generated using ChromoMap (Anand and Rodriguez Lopez, 2022) in RStudio.

### Low numbers of *Theobroma cacao* NLR-type genes compared to other tree species

Resistance genes of the NLR-type are numerous and highly polymorphic both within and between plant species (Van de Weyer *et al*., 2019). Trees have generally been shown to have higher numbers of NLR-type genes than herbaceous species, and this is suggested to accomodate a changing pathogen environment over the life of a tree (Tobias and Guest, 2014). More recently, analyses of gene content in diploid genomes have been possible due to advances in sequencing and computational technologies. Notably, tree resistance genes have been investigated within the diploid genome for *Melaleuca quinquenervia* (Chen *et al*., 2023), highlighting variation in numbers (763 and 733) between haplotypes and across chromosomes for this class of gene. Our analysis of the NLR complement within *T. cacao* showed similar representation to an earlier genome study (Argout *et al*., 2011) that determined just 297 non-redundant NLR-type genes. Our work, however, presents the diploid content of NLRs as 293 and 326 for haplotype A and B respectively, permitting deeper investigation of homologs at loci. We confirmed low numbers and percentages of the Toll/Interleukin-1 receptor (TIR) NLR class (termed TNLs) of resistance genes within *T. cacao* (Figure 3) compared to many trees within the Rosid clade, in which they comprise 40 - 60% (Arya *et al*., 2014; Christie *et al*., 2016; Kohler *et al*., 2008). This was noted in earlier molecular and genomic studies where only 4% of NLRs contained the TIR domain (Argout *et al*., 2011; Kuhn *et al*., 2006), with our classification indicating 3.8% and 5% for haplotype A and B. Unlike the previous study with primers for molecular markers that found TNLs on chromosome three and five (Kuhn *et al*., 2006), we determined TNLs within clusters on chromosome 4 (three genes), 5 (two genes) and 6 (three and five genes on haplotype A and B), plus a single TNL gene on chromosome 2, 4 and 6 of each haplotype. Our finding of only five and 12 NLR-type genes with non-canonical integrated domains in the two haplotypes also suggests that this integration is less important as a strategy for pathogen recognition and response in cultivated *T. cacao*, compared to other species (Marchal *et al*., 2022).

**Figure 3.**
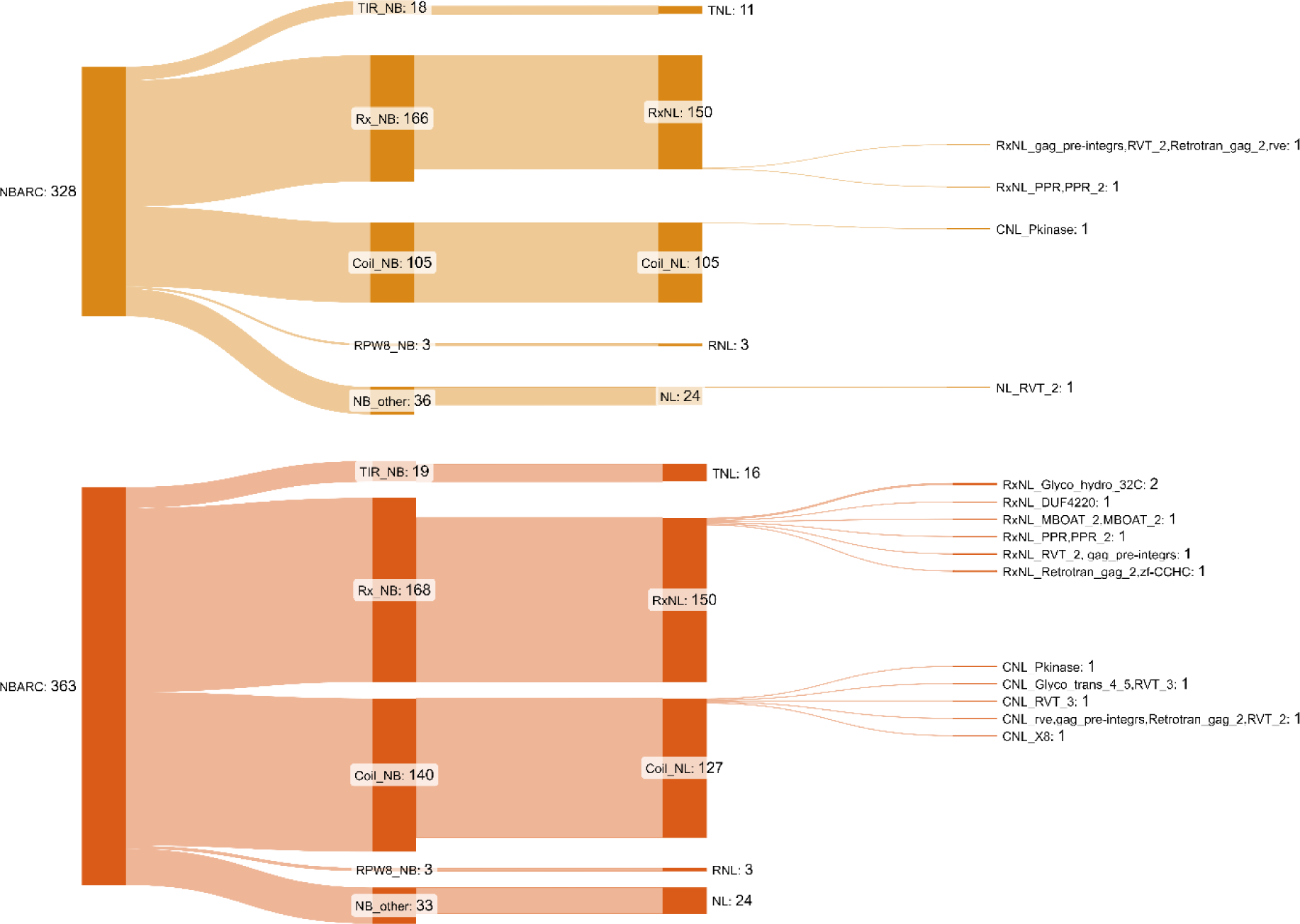
**The predicted NLR gene complement in the parentally phased *Theobroma cacao* genome. The two sets of chromosomes corresponding to (above) Haplotypes A and (below) B were independently classified** (Chen *et al*., 2023) **and visualised to present the domain classes using SankeyMatic (Bogart, 2014) including novel integrated domains (IDs) with abbreviations derived from Pfam database** (Mistry *et al*., 2021). **NBARC = Nucleotide Binding Domain, TIR = Toll/Interleukin-1 receptor, Rx = Potato CC-NB-LRR protein Rx, CC=Coil-Coil Domain, RPW8 = RESISTANCE TO POWDERY MILDEW 8-like coiled-coil**

### *Theobroma cacao* NLR-type genes are present and polymorphic at QTLs for VSD resistance

Informative QTL regions were investigated for resistance gene homologs across the parental inherited chromosomes. The most closely aligned VSD QTLs between PNG and Sulawesi *T. cacao* progeny trials were on chromosomes 3 and 9, at around 29 and 31 Mb, while chromosome 8 harboured a region between 6 – 7 Mb (Table 3, experimental procedures). We checked these regions based on our annotations for NLRs and identified an NLR cluster of two predicted genes at around 29 Mb on **chromosome 3** within both parentally inherited alleles. The location matches almost exactly with our calculated region based on the reversed start position for the cM QTL (Table 3). When we aligned the four amino acid sequences for these NLRs with ClustalW (Thompson *et al*., 1994) and visualised in BioEdit (Hall, 1991), we see close homology across the cluster but variation in one haplotype A protein, notably an insertion and deletion in the sequence for gene identity g2391.t1 (Figure 4, above). We found an NLR cluster of three genes on **chromosome 8** at around 3.8 Mb which falls within the 2 cM interval specified by Singh (2021) of 2 - 4 Mb from the QTL locus midpoint. The NLR cluster formed two orthogroups, one group of two head-to-head Rx-NLR predicted proteins from each haplotype (g1318.t1, g1320.t1 and g3681.t1, g3683.t1) and the other of single, nearly exactly homologous proteins, with one amino acid variant, namely proline (g1316.t1, Haplotype A) to arginine (g3679.t1, Haplotype B). Between the region 27 – 36 kb on **chromosome 9** we determined four NLRs from Haplotype A and three from Haplotype B. Paired homologs closest to the calculated QTL mid-point location at around 31 Mb on chromosome 9 were aligned and show an insertion/deletion as well as five single amino acid substitutions (Figure 4, below).The single predicted Rx-NLR gene on chromosome 9 of Haplotype A at 27 Mb has no homolog in Haplotype B.

**Figure 4.**
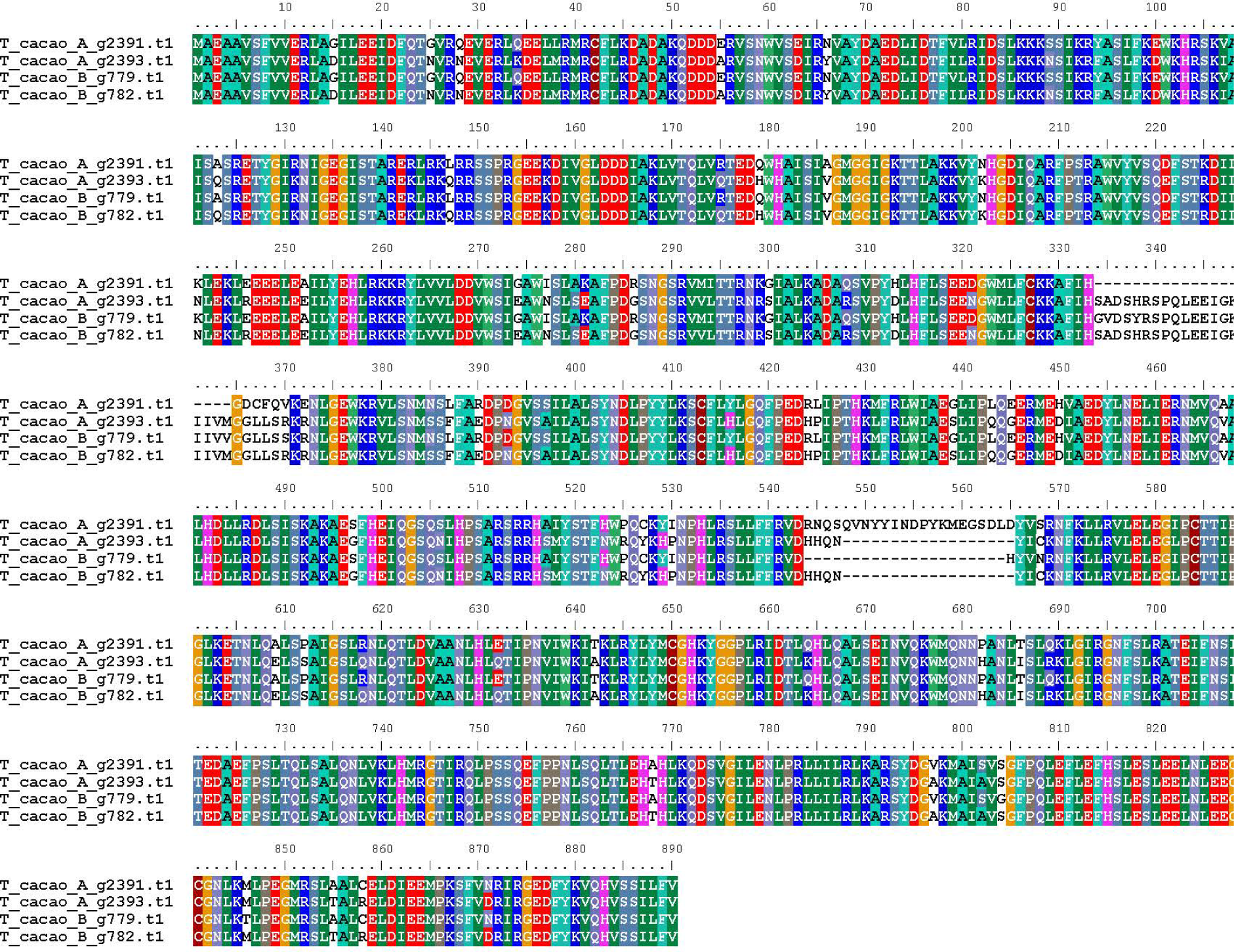
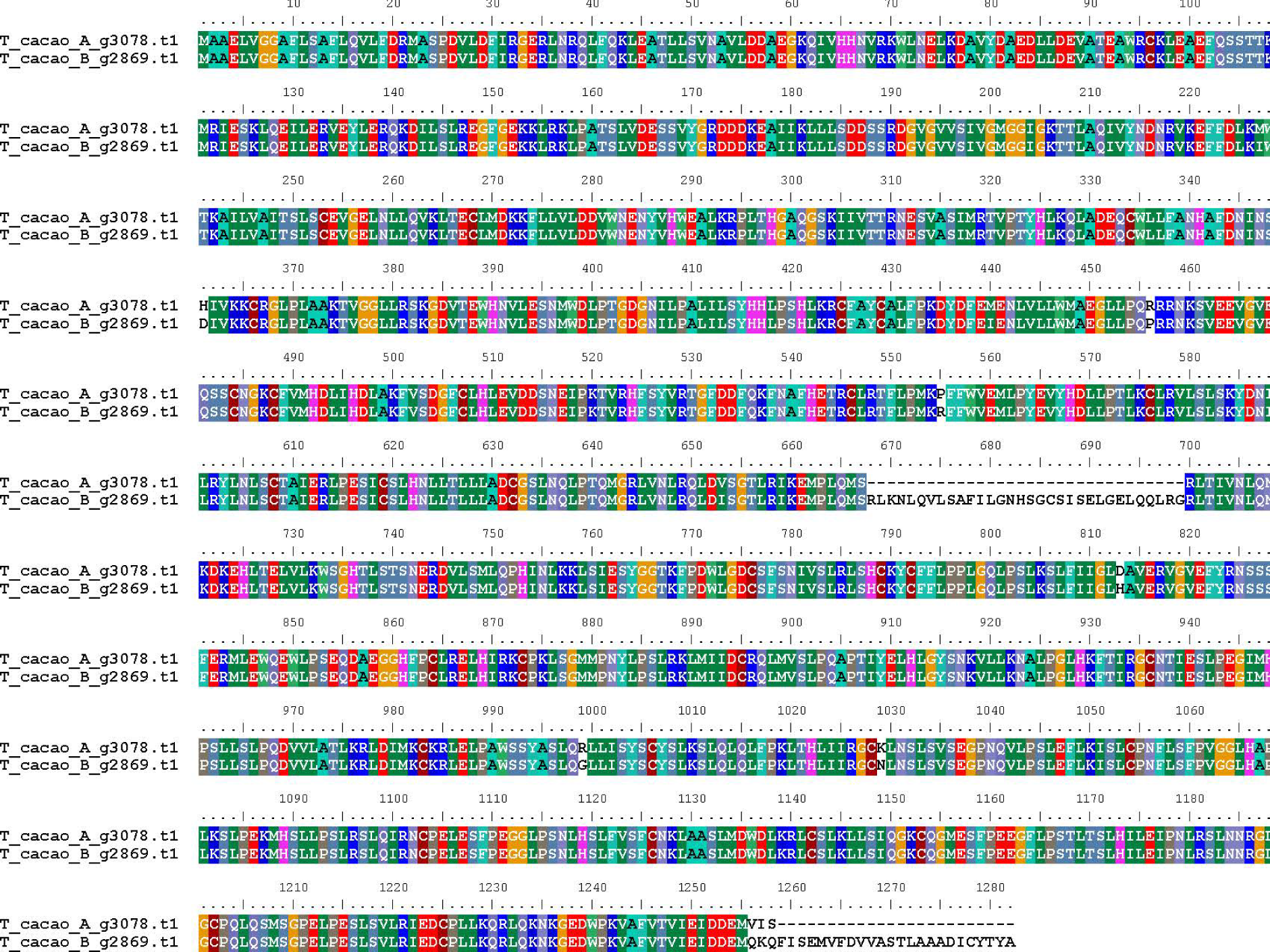
**Amino acid sequence alignment of an NLR cluster matching the QTL mapped regions at around 29 Mb on chromosome 3 (above) and single NLR homologs at 31 Mb chromosome 9 (below) within both parentally inherited alleles, A and B, for *Theobroma cacao* clone 26. All predicted NLRs are annotated as Rx-NLRs** (Chen *et al*., 2023) **named for resistance to potato virus X.**

The QTLs on chromosome 3 and 8 accounted for 11 and 15% phenotypic variance in the Sulawesi progeny trials and were informative for VSD and PPR resistance in the PNG trial (Table 3). The chromosome 9 QTL accounted for 16 - 18% phenotypic variance in the Sulawesi trial. All putative resistance genes at QTLs were annotated as Rx NLR-type resistance genes. The Rx-NLR coded protein from potato (*Solanum tuberosum*) has been identified as conferring resistance to the potato virus X by direct recognition. Recent work has clarified the interaction of sensor Rx-NLRs with downstream helper NLR protein oligomerisation (Contreras *et al*., 2023), showing a distinct mechanism for immune responses in plants. While our work has uncovered the arrangement and sequence differences of predicted NLR clusters at QTLs for resistance to VSD, confirmation of these genes is needed across multiple progenies that have clear phenotypes for resistance/susceptibility. Amplification of DNA regions around these clusters will provide additional clarity but was beyond the scope of the current study. Functional validation would also be beneficial to determine mechanisms of recognition and response as it is known that paired head to head NLRs, as we identified on chromosome 8, can determine resistance to more than one pathogen (Xi *et al*., 2022), potentially explaining the results from the study conducted in PNG.

### Annotation statistics show high homology between allelic pairs in *Theobroma cacao*

The orthology of all annotated proteins was analysed with OrthoVenn3 as described previously. The predicted proteins indicated 22,486 shared clusters, of which 21,800 included a single predicted homologous gene from each haplotype. The largest cluster had 44 predicted proteins and there were 9.76% singletons. In total there are 2,928 (2,896), 1,865 (1,798), 3,278 (3,315) predicted genes for Haplotype A (and B) on chromosomes 3, 8 and 9 respectively. Except for chromosome 9, with around 76 predicted genes within ∼2 Mb of the QTL informing resistance to VSD, the locations on chromosomes 3 and 8 are in highly dense gene-encoding regions, with 325 and 370 predicted genes. We investigated the predicted genes at the QTLs and found that homologs were present between haplotypes (Table 2) and we reviewed their predicted protein functions based on Pfam database (Mistry *et al*., 2021) annotations. Due to the density of predicted genes, we limited our investigations to immediate proximity around NLRs that corresponded to QTLs. Around the predicted QTL for resistance to VSD on chromosome 3, the Rx-NLR gene cluster is interspersed with two predicted genes coding for protein kinases (Table 2). The kinases have fused LRR domains and predicted transmembrane motifs indicating extracellular receptors and a putative role in defence response. The importance of this class of receptor-like kinase (RLK) has been understood in rice with studies showing *Xa21* confers resistance to *Xanthomonas* sp. and multiple other pathogens (Ercoli *et al*., 2022). The finding of kinases so closely associated with the Rx-NLR genes on chromosome 3 in both parental alleles, at the predicted region for VSD resistance QTL, suggests a potential role in disease response and a target for further research. The NLR QTL regions associated with resistance on chromosomes 8 and 9 did not have associated disease response-like genes, however it is likely that multiple genes are involved in the VSD resistance response observed.

**Table 2.** Predicted and annotated genes that are in proximity to the QTLs for resistance to *Ceratobasium theobromae*, causal agent of vascular streak dieback in cultivated *Theobroma cacao*. The annotated genomes are available at NCBI: PRJNA1083235 and PRJNA1086984.

| Haplotype A | Haplotype B | Pfam ID | Annotation |
| --- | --- | --- | --- |
| NLR QTL chromosome 3 (40 kb region) |  |  |  |
| V9T21_008377 | WDX93_008321 | PF00334 | Nucleoside diphosphate kinase |
| V9T21_008378 | WDX93_008322 | PF01625 | Peptide methionine sulfoxide reductase |
| V9T21_008380 | WDX93_008324 | PF07714 | Protein tyrosine and serine/threonine kinase |
| V9T21_008380 | WDX93_008324 | PF13855 | Leucine rich repeat |
| V9T21_008382 | WDX93_008326 | PF07714 | Protein tyrosine and serine/threonine kinase |
| V9T21_008382 | WDX93_008326 | PF13855 | Leucine rich repeat |
| V9T21_008382 | WDX93_008326 | PF08263 | Leucine rich repeat N-terminal domain |
| V9T21_008383 | WDX93_008327 | PF02298 | Plastocyanin-like domain |
| NLR QTL chromosome 8 (10 kb region) |  |  |  |
| V9T21_020396 | WDX93_020310 | PF00745 | Glutamyl-tRNA<sup>Glu</sup> reductase, dimerisation |
| V9T21_020396 | WDX93_020310 | PF01488 | Shikimate / quinate 5-dehydrogenase |
| V9T21_020397 | WDX93_020311 | PF01293 | Phosphoenolpyruvate carboxykinase |
| V9T21_020399 | WDX93_020313 | PF08694 | Ubiquitin-fold modifier-conjugating enzyme1 |
| V9T21_020401 | WDX93_020315 | PF12295 | Symplekin tight junction protein C terminal |
| V9T21_020401 | WDX93_020315 | PF11935 | Symplekin/PTA1 N-terminal |
| V9T21_020402 | WDX93_020316 | PF11935 | Symplekin/PTA1 N-terminal |
| V9T21_020404 | WDX93_020318 | PF00719 | Inorganic pyrophosphatase |
| NLR QTL chromosome 9 (3 kb region) |  |  |  |
| V9T21_023576 | WDX93_023431 | PF00514 | Armadillo/beta-catenin-like repeat |
| V9T21_023578 | WDX93_023433 | PF02341 | RbcX protein |
| V9T21_023579 | WDX93_023434 | PF00076 | RNA recognition motif |

### Concluding remarks

A recent study, using single nucleotide polymporphisms, confirmed low genetic diversity due to selection for domestication in cultivated cacao (Orduña *et al*., 2024). The results of that work may explain why similar loci are apparent across varieties for resistance to different diseases, as was found in the two studies, in Indonesia and in PNG, that informed the current research. Genome resources have, until recently, been presented as collapsed haploid genomes and the allocation of QTLs has been limited due to loss of valuable information from the two allele sets. Technologies are rapidly improving and permitting far more detailed comparative studies than were possible only a few years ago. Our original goal of assembling both the host and compatible pathogen genomes from one sample was unsuccessful due to low sequence coverage, only 1X HiFi reads, for *C. theobromae* reads. However future methods could use adaptive sequencing with the Oxford Nanopore Technologies metagenome sampling tools (Martin *et al*., 2022). This approach is rapidly developing, with new models incorporating Bayesian dynamic adaptive sampling (Weilguny *et al*., 2023), and providing promising tools to investigate the biology of unculturable biotrophic pathogens.

For this research we employed recent advances in technology and software to assemble and assign parental chromosomes for a VSD susceptible *T. cacao* genotype (clone 26) from the Mars Cocoa Research Centre in Pangkep, Sulawesi, Indonesia. We annotated the two parental chromosome sets independently and specifically we characterised all the predicted NLR-type resistance genes. The resulting curated and annotated genomes were compared at QTL locations that were identified to explain resistant phenotypes to VSD and PPR. Our results show that the QTLs may indeed resolve to NLR-type resistance genes, and that these may be important to investigate further for breeding trees with resistance to VSD, PPR, and potentially other diseases of cacao. Our work provides a more comprehensive analysis of the predicted NLRs and other genes on each chromosome and shows the variation in structure and type of these genes within a VSD susceptible individual.

Future work could develop molecular screening around these specific genes across both resistant and susceptible trees for a range of diseases of cacao. The current work has developed foundational genome resources to facilitate such analyses that may be used to assist future breeding programs for resistance.

### Experimental procedures

#### High Molecular Weight DNA extraction protocol testing

Prior to our work in Sulawesi, Indonesia, we ran exhaustive high molecular weight (HMW) DNA extraction protocol testing on cacao leaf samples from plants maintained in the University of Sydney greenhouses. A persistent problem remained for clean extractions due to high contaminant levels that had the properties of a jelly-like substance, presumably pectin, though not confirmed. We attempted to resolve the nature of the contaminant with enzymatic testing using pectinase. We also tested our contaminant DNA with nuclear magnetic resonance spectrometer (NMR). Data was collected on a 600 MHz spectrometer equipped with triple resonance CryopProbe and the following buffer conditions: 10 mM Tris; 10% D2O; pH 8.0, in 3 mm short NMR tubes (volume ∼160 µl). We ran controls of 10 mM Tris buffer and clean HMW DNA in Tris buffer. Test samples included pectin, pectin with pectinase treatment, contaminated DNA with pectinase treatment, contaminated DNA with macerozyme (incl. pectinase) treatment. Unfortunately the signals in the aromatic/amide regions were beyond detection, suggesting a large DNA-contaminant (potentially covalent) complex form in this sample. Young leaves were found to produce cleaner DNA but did not show any symptoms of VSD, so we aimed to extract samples from older infected leaves using a combined protocol described below that improved on but did not resolve the contaminant problem.

#### Cacao leaf material collected from field trial tree

We collected fully expanded leaf material on the 10 January 2023, during wet weather, from the Mars Cocoa Research Institute field trials in Pangkep, South Sulawesi, Indonesia. The leaves were found on T. *cacao* F1 generation progeny, VSD susceptible clone number 26, from Sulawesi 1 – S1 (resistant maternal), crossed with RUQ 1347 (MIS_GBR207 CCN 51) (susceptible paternal) parental plants (Turnbull et al., 2017). The leaves were symptomatic of VSD with browning of the leaf apex and vascular death visible in the petiole (Figure 5). Leaves were placed in plastic bags and transported to the laboratories at Universitas Hasannudin, Makassar and refrigerated overnight.

**Figure 5.**
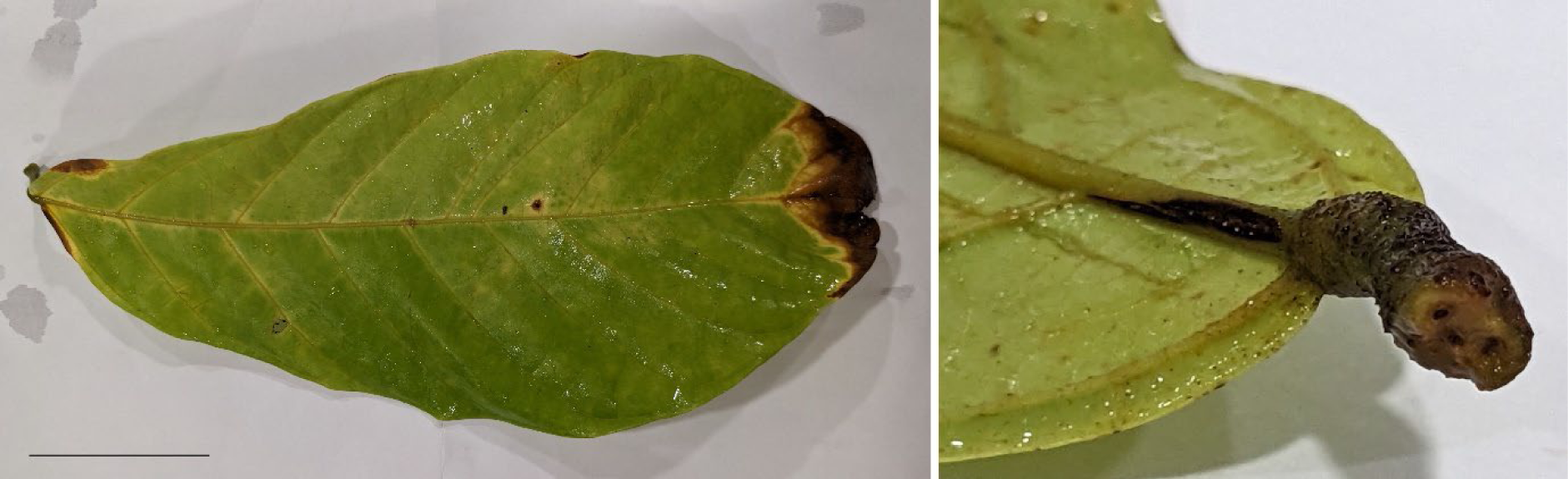
Leaf from *Theobroma cacao*, VSD susceptible clone 26, growing at the Mars Cocoa Research Institute field trials, Pangkep South Sulawesi, Indonesia. (Left) The symptomatic leaf apex dieback and some chlorosis indicating *Ceratobasium theobromae*. (Right) Visible vascular death caused by the pathogen. Scale bar left ∼5 cm.

#### High Molecular Weight DNA extracted from cacao leaves

The following day while leaves were fresh, HMW DNA was extracted using a protocol adapted from the sorbitol wash method developed by Jones et al. (2021), and the HMW plant DNA CTAB extraction method developed by Hilario (2018). Small (1 cm^2^) pieces of the leaves were placed into 1.5 mL centrifuge tubes with a small amount (a few mg) of PVP40000. The tissue was flash frozen in liquid nitrogen and ground into a powder with micropestle while frozen. Three or more rounds of sorbitol wash were used on the samples, in accordance with the protocol (Jones et al., 2021), until the supernatant was no longer viscous. Then DNA was extracted, according to the protocol (Hilario, 2018), using CTAB buffer with an incubation step at 56 °C for 2 hours, in a water bath. Apart from an extended RNase A incubation step, of 10 mins, the protocol was followed until the precipitation step. The DNA formed a visible gelatinous precipitate at this stage and was left to continue precipitating at −20 °C overnight. The following day the DNA was collected by centrifugation at 3000 rpm for 25 min at room temperature and cleanup steps followed according to the protocol before dissolving in 10 mM Tris-HCL pH 8.0. The DNA was quantified, and purity was checked using a NanoDrop™ Lite Plus spectrophotometer and a QuBit™ 2.0 Fluorometer. Purified HMW DNA (10 x 1.5 µL tubes) was couriered under an Australian Department of Agriculture, Water, and the Environment import permit to the Australian Genome Research Facility (AGRF), Brisbane, Australia, for cleanup, size selection, library prep and sequencing on Pacific Biosciences of California, Inc. (PacBio) Sequel II.

#### Formaldehyde cross-linking of leaves for HiC sequencing

We prepared samples according to the manufacturer’s protocols for chromatin crosslinking library prep and sequencing at Phase Genomics (Seattle, Washington, US) to obtain HiC reads. Five grams of leaf material was immersed in 2% (v/v) sodium hypochlorite for 2 mins for surface sterilisation and then washed with ultrapure water. The leaves were cut into strips and immersed in 1% (v/v) formaldehyde for 40 minutes with intermittent mixing. The crosslinking reaction was quenched with the addition of glycine to a final concentration of 125 mM for 15 minutes. The crosslinked leaves were washed with ultrapure water, dried, then ground to a powder after freezing with liquid nitrogen.

#### Data processing and genome assembly

Details of all the software, versions, parameters and scripts used to process the data are available here: https://github.com/peritob/Theobroma-cacao-genome/tree/main

#### Preparing the raw data for processing

We downloaded the 23 GB (ccs.bam) HiFi sequence read data from AGRF, filtered the reads using HiFiAdapterFilt (Sim *et al*., 2022) and checked the statistics using Nanoplot (De Coster and Rademakers, 2023). The HiC paired-end read (25 GB) mapping statistics against the cacao Criollo reference genome (GCF_000208745.1) were good for informative read pairs (12.18%) but not against the pathogen genome (GCA_012932095.1) at (0.03%), although 4.28% same strand high quality reads were mapped. We mapped all the filtered HiFi reads to the *C*. *theobromae* genome (Ali *et al*., 2019) with Minimap2 (Li, 2018) to filter the pathogen from host. We presumed all the mapped reads to be pathogen and all the unmapped reads to be from the host and proceeded accordingly. Additionally, we ran a local blast with internal transcribed spacer (ITS) primers on the VSD mapped raw sequence reads. This confirmed a single exact *in silico* hit using the ITS regions using forward primers ITS4 (White *et al*., 1990) and reverse ITS5A (Stanford *et al*., 2000). Raw Illumina data available for the parental plants was trimmed of adaptors and low quality reads with Fastp (Chen *et al*., 2018).

#### Genome assembly and chromosome scaffolding

We attempted several assembly approaches with Hifiasm software (Cheng *et al*., 2021). Previous experience with HiC scaffolding has shown the possibility to phase the separate nuclear compartments (Tobias *et al*., 2022). We therefore attempted to build a plant-pathogen hybrid genome to scaffold with HiC and visually identify and separate the two organisms using the heatmaps. The hybrid genome did not complete, likely due to insufficient overlaps for the pathogen reads. We tried to assemble all the pathogen reads into consensus sequences using Flye (Kolmogorov *et al*., 2019) and Hifiasm without success. We then tested the trio-binning option for Hifiasm of the diploid *T. cacao* incorporating the Illumina parental reads after running Yak (Li, 2020). This was unsuccessful, so we assembled the diploid *T. cacao* genome using HiC integration with Hifiasm followed by HiC scaffolding of the resulting genomes independently with Juicer (Neva C. Durand *et al*., 2016), 3D-DNA (Dudchenko *et al*., 2017) and manual scaffolding with Juicebox (Neva C Durand *et al*., 2016).

### Theobroma cacao genome curation

We used the Matina 1-6 cacao genome (GCF_000403535.1) to anchor our genome scaffolds with PAFScaff (Field *et al*., 2020) based on Minimap2 (Li, 2018) mapping. We screened the resulting fasta files for non-nuclear genomic contaminants with FCS-GX (Astashyn *et al*., 2024) and Tiara (Karlicki *et al*., 2022). In each case, the two haplotypes were run independently. FCS-GX was run against the NCBI gxdb (build date 2023-01-24; downloaded 2023-09-12). Additional checks of contamination were executed with Taxolotl (Tobias *et al*., 2022). Taxolotl was provided with the annotation for each haplotype and run with taxwarnrank=class using mmseqs2 against the NCBI nr database (compiled 2022-05-19). In addition to generating results for all sequences in each haplotype, analysis of subsets was performed for (1) chromosome scaffolds only, (2) pure FCS-GX scaffolds, (3) contaminated FCS-GX scaffolds, (4) Tiara “eukarya” scaffolds, and (5) scaffolds not classified as “eukarya” by Tiara. Results were visualised with Pavian. Lists were made of non-plant contaminant contigs from the fcs_gx_reports and were removed, as well as plastid contigs flagged in the tiara outputs. We then ran Telociraptor (Edwards, 2023) followed by Chromsyn (Edwards *et al*., 2022) on the chromosome-level contigs to compare synteny and scaffolds tweaked according to the contig graphs and figure outputs. We used the parental data available within our lab (Singh, 2021) to assign scaffolds and contigs. We mapped the maternal S1 data to the concatenated haplotype A and B genomes using Hisat2 (Kim *et al*., 2019) and then similarly mapped the RUQ 1347 data. We assumed that the most reads mapped were the likely parental chromosome/contig and curated each genome. Of the chromosomes, 3, 5 and 8 were switched accordingly (Figure 6). We designated the S1 maternally inherited contigs as genome T_cacao_v1.3_A, and RUQ 1347 paternally inherited as T_cacao_v1.3_B.

**Figure 6.**
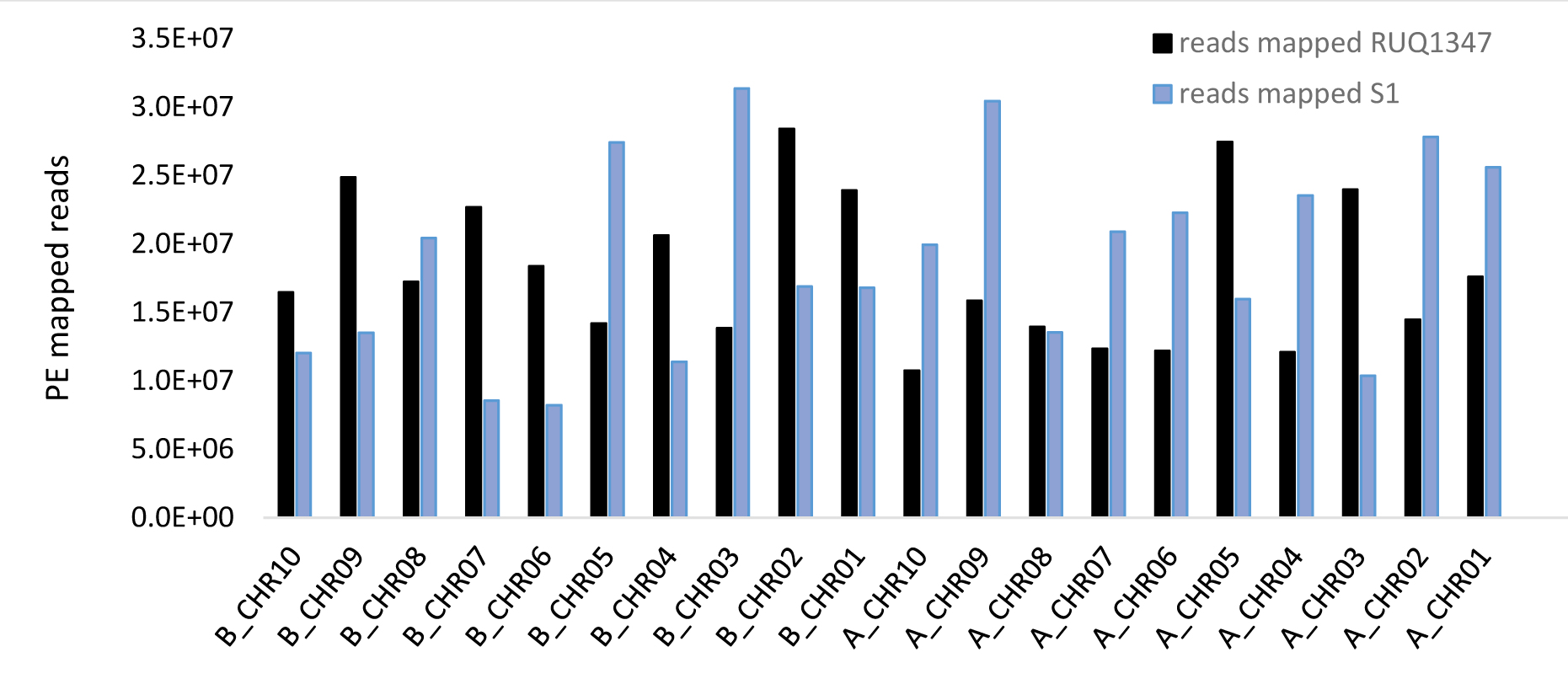
*Theobroma cacao* resistant (S1) and susceptible (RUQ 1347) parental paired-end (PE) Illumina sequence reads mapped to the concatenated Hap A and B progeny chromosomes. The reads that were mapped in greater numbers to chromosomes and contigs were used to determine parental assignment and swapped accordingly.

#### Parental assigned genomes were annotated independently

To repeat mask the genome for annotation we ran both genomes through Repeatmasker and Repeatmodeler. We then mapped publicly available RNAseq data from CCN51 *Theobroma cacao* from NCBI (PRJNA933172) to the soft-masked genomes independently with hisat2. We trimmed the RNAseq data with Fastp (Chen *et al*., 2018) prior to mapping and processed the outputs with Samtools (Li *et al*., 2009) to make bam files. The mapped RNAseq bam files were used as evidence to predict protein coding genes within the funannotate (Palmer and Stajich, 2023) *predict* pipeline with the BUSCO embryophyte (Manni *et al*., 2021; Simão *et al*., 2015) database. We used Interproscan5 (Jones *et al*., 2014) independently and then ran funannotate *annotate* to obtain final predicted proteins and coding sequences. In order to make GenBank (Sayers *et al*., 2023) compatible annotations, we ran the resulting files with Table2asn (GenBank, 2023).

#### NLR-type resistance gene annotation

We annotated the resistance gene complement with the FindPlantNLRs pipeline (Chen *et al*., 2023), that incorporates NLR-Annotator (Steuernagel *et al*., 2020), HMMER (Finn *et al*., 2015; Johnson *et al*., 2010) and BLAST (Altschul *et al*., 1990) components as well as Interproscan5 (Jones *et al*., 2014). Unlike many annotation methods, the pipeline takes an unmasked genome as a starting point to avoid masking, and hence not annotating, repetitive LRR regions that are characteristic of NLR-type genes. The last stage of the pipeline determines NLR gene classes, including predicted integrated domains, and locations within each parental haplotype. All NLRs were plotted to visualise on chromosomes with Chromomap (Anand and Rodriguez Lopez, 2022) and classes, based on domain Pfams, were visualised with Sankeymatic (Bogart, 2014).

#### QTL locations on chromosomes predicted and compared against NLR-type gene predictions

Informative QTLs for resistance to VSD were obtained from data of two previous studies (Epaina, 2012; Singh, 2021). Using the chromosome lengths for our parental genome haplotypes we calculated the predicted locations for VSD resistance QTLs. In brief, we took averages of Linkage Group (LG) map lengths (Epaina, 2012), and divided the chromosome base pair length from assembled genomes (current study) by centimorgan (cM). We then determined the expected mid-point (in base pairs) for the resistance QTLs. Singh (2021) predicted QTLs confidence intervals at 3-4 Mb and determined three significant QTLs; on chromosome 3, 8 and 9. Initially our calculations for the QTL locations from the PNG progeny study showed a significant region at around 18.5 Mb on chromosome 8 (Table 3). This was not concordant with the Sulawesi study, at between 5.5 and 8.8 Mb, leading us to investigate if our calculations were misrepresenting the ‘start’ position for chromosome eight. When recalculated based on the reverse orientation, we found that the QTL matched more closely, at around 7 Mb. We also noted that the QTL identified on chromosome 3, when start positions were reversed, indicated a cluster of NLR-type genes at around 30 Kb. Several significant QTLs were associated with resistance to other pathogens including PPR (Table 3) in the study by Epaina (2012). QTLs that were in similar regions within both studies were regarded as likely harbouring useful candidate genes for breeding resistance. The most closely aligned VSD QTLs between studies were on chromosomes 3, 8 and 9. Detailed comparisons for the VSD significant QTLs across the two studies were undertaken to investigate resistance gene homologs and other predicted genes across the parental chromosomes.

**Table 3.** QTLs for vascular streak dieback (VSD) resistance were calculated using average *Theobroma cacao* Linkage Group (LG) lengths (Epaina, 2012) and dividing chromosome lengths (bp) by LG lengths (in determine 1 cM in bp (per chromosome). QTLs (cM) were then multiplied by bp length for 1 cM to obtain the chromosome locations for the diploid genome. Green cells and brown cells are VSD and *hthora* pod rot (PPR) resistance QTL physical locations on chromosomes.

| Theobroma cacao A (S1 inherited) |  |  |  |  |  |  |  |  |  |  |  |
| --- | --- | --- | --- | --- | --- | --- | --- | --- | --- | --- | --- |
| Chromosomes (LG) |  | 1 | 2 | 3 | 4 | 5 | 6 | 7 | 8 | 9 | 10 |
| cM (average) |  | 99.9 | 107.2 | 91.6 | 79.7 | 89.8 | 75.8 | 56.6 | 60 | 106.1 | 62.4 |
| bp per chromosome (this study) |  | 39,005,981 | 41,965,441 | 38,945,508 | 33,857,508 | 40,959,067 | 25,938,454 | 26,721,266 | 25,579,687 | 40,159,845 | 25,091,101 |
| bp per cM |  | 390,450 | 391,469 | 425,169 | 424,812 | 456,114 | 342,196 | 472,107 | 426,328 | 378,509 | 402,101 |
| Singh (2021) | VSD QTL (bp) |  |  | 9,353,725<br>(*29,591,783) |  |  |  |  | 8,782,359 | 25,170,874 |  |
| Epaina (2012) | VSD QTL (bp) | 11,270,737 | 26,861,405 | 8,531,872<br>(*30,413,636) | 27,163,747 | 7,976,071 |  |  | 18,514,151<br>(*7,065,536) | 31,452,994 |  |
| Epaina (2012) | PPR QTL (bp) |  |  | 8,531,872<br>(*30,413,636) | 23,158,620 | 6,937,499 |  |  | 18,514,151<br>(*7065,536) |  |  |
| Theobroma cacao B (RUQ 1347 inherited) |  |  |  |  |  |  |  |  |  |  |  |
| Chromosomes (LG) |  | 1 | 2 | 3 | 4 | 5 | 6 | 7 | 8 | 9 | 10 |
| bp per chromosome (this study) |  | 39,156,173 | 42,761,000 | 39,170,102 | 34,430,140 | 41,360,500 | 29,284,707 | 25,927,970 | 21,951,902 | 41,168,245 | 24,929,739 |
| bp per cM |  | 391,954 | 398,890 | 427,621 | 431,997 | 460,585 | 386,342 | 458,091 | 365,865 | 388,014 | 399,515 |
| Singh (2021) | VSD QTL (bp) |  |  | 9,407,666<br>(*29,762,436) |  |  |  |  | 7,536,820 | 25,802,906 |  |
| Epaina (2012) | VSD QTL (bp) | 11,314,135 | 27,370,630 | 8,581,075<br>(*30,589,027) | 27,623,167 | 8,054,243 |  |  | 15,888,421<br>(*6,063,481) | 32,242,768 |  |
| Epaina (2012) | PPR QTL (bp) |  |  | 8,581,075<br>(*30,589,027) | 23,550,302 | 7,005,492 |  |  | 15,888,421<br>(*6,063,481) |  |  |

\*QTL determined with start position on chromosome reversed.

## Acknowledgements

The Joint Cocoa Research Fund, European Cocoa Association supported this research. The authors acknowledge the facilities, and the scientific and technical assistance of the Sydney Informatics Hub at the University of Sydney and, in particular, access to the high-performance computing facility Artemis. We acknowledge the support of Sydney Analytical for running samples and interpretation on the University of Sydney Nuclear Magnetic Resonance (NMR) Core Facilities. Thank you to Mark Powrie who generously assisted during the work in Sulawesi.

## Author contributions

DG and PT initiated and led the project. PT and JD collected the material from established clonal trials developed and managed by AP. JM and AP supported and facilitated the research in Sulawesi. JD, PT, AZ, EB extracted HMW DNA and cross-linked samples. PT and EB organised sample deliveries. PE and GS provided results of QTL trials that were supported by DG and RP. GS provided parental data. PT assembled, curated, annotated the genomes, and interpreted results, RE ran contamination screening of genomes. PT submitted the data to NCBI and wrote the manuscript. All authors contributed comments and edits to the manuscript.

## Data availability statement

All data that support the findings of this research are publicly available. Raw data is available at National Centre for Biotechnology Information (NCBI) under the following biosample accession: SAMN40241783 and bioproject accession: PRJNA1083235. Annotated diploid *Theobromae cacao* (clone 26) genomes are available here: PRJNA1083235 (maternally inherited), PRJNA1086984 (paternally inherited). NLR predicted coding and amino acid sequences and gff3 data is available at https://github.com/peritob/Theobroma-cacao-genome/tree/main.

## Abbreviations

NLR: – nucleotide binding site leucine-rich repeat gene models
LRR: – leucine-rich repeat domain
GB: – gigabyte
Gb: – Gigabase
Mb: – Megabase
kb: – kilobase
VSD: – vascular streak dieback, causal agent *Ceratobasidium theobromae*
PPR: – Phytophthora Pod Rot, causal agent *Phytophthora palmivora*

